# Allele-specific epigenetic regulation of *FURIN* expression at a coronary artery disease susceptibility locus

**DOI:** 10.1101/2023.05.19.541547

**Authors:** Wei Yang, Junjun Cao, David G. McVey, Shu Ye

## Abstract

**Background:** Genome-wide association studies have revealed an association between the genetic variant rs17514846 on chromosome 15q26.1 and coronary artery disease susceptibility. Studies have shown that rs17514846 influences the expression of the FES Upstream Region (*FURIN*) gene located at this locus in monocytes. We investigated the mechanism through which rs17514846 modulates *FURIN* expression.

**Methods and Results:** An analysis of isogenic monocytic cell lines with either the rs17514846 C/C or A/A genotype showed that the cells of the A/A genotype expressed higher levels of *FURIN* than cells of the C/C genotype. A pyrosequencing methylation analysis showed that the cytosine (in a CpG motif) at the rs17514846 position on the C allele was methylated. Treatment with the DNA methylation inhibitor 5-aza-2’-deoxycytidine increased *FURIN* expression. A bioinformatic analysis indicated that the rs17514846 site might interact with the transcription factor MeCP2 that often functions as a gene repressor by binding to methylated CpG sites. An electrophoretic mobility super-shift assay with a probe corresponding to the DNA sequence at and around the rs17514846 position of the C allele detected two DNA-protein complex bands, which were altered by the addition of an anti-MeCP2 antibody in the assay, whilst these DNA-protein complexes were barely detectable with a probe for the A allele. A chromatin immunoprecipitation assay showed an enrichment of the DNA sequence containing the rs17514846 site in chromatin precipitates pulled down by an anti-MeCP2 antibody. siRNA-mediated knockdown of MeCP2 caused an increase in *FURIN* expression. Furthermore, MeCP2 knockdown increased monocyte migration and proliferation, and this effect was diminished by a FURIN inhibitor.

**Conclusions:** The results of our study suggest that DNA methylation inhibits *FURIN* expression and that the coronary artery disease predisposing variant rs17514846 modulates *FURIN* expression and monocyte migration via an allele-specific effect on DNA methylation.

## Introduction

Genome-wide association studies have revealed a robust association between genetic variation on chromosome 15q26.1 and coronary artery disease (CAD) susceptibility^1–4^. The A allele of the lead single nucleotide polymorphism (SNP), rs17514846, at this locus is associated with increased CAD risk^1–4^. rs17514846 is located in a noncoding region of the *FURIN* (<u>F</u>ES <u>U</u>pstream <u>R</u>egio<u>n</u>) gene and modulates monocyte *FURIN* expression^5^.

The *FURIN* gene encodes the subtilisin-like proprotein convertase FURIN that possesses proteolytic activity to cleave the prodomain off of, and thereby activates, many proteins. There is accumulating evidence indicating that FURIN plays an important role in atherosclerosis, the pathological condition underlying CAD. A key process in the development and progression of atherosclerosis is the recruitment of monocytes into the arterial wall and atherosclerotic lesions^6^. Our recent study showed that FURIN promotes human monocyte migration and proliferation^5^. In agreement, a mouse model study by other investigators demonstrated that FURIN inhibition reduces monocyte migration and retards atherosclerotic lesion progression^7^. Furthermore, a clinical study reported an association of high FURIN levels with increased recurrent cardiovascular events and mortality in patients with acute myocardial infarction^8^.

It is currently unknown as to how the CAD-associated SNP rs17514846 modulates *FURIN* expression. Our present study sought to investigate the underlying regulatory mechanism. The study finds that the non-risk allele (the C allele) is subjected to methylation of a CpG motif at the rs17514846 site, interacts with the transcription factor MeCP2 (that often acts as a gene repressor by binding to methylated CpG sites^9,10^) and expresses less *FURIN*, whereas the C to A change of this SNP in the CAD risk allele (the A allele) abolishes the CpG motif, diminishes MeCP2 binding and leads to increased *FURIN* expression. Furthermore, our study shows that targeted knockdown of MeCP2 results in an increase of *FURIN* expression and promotes monocyte migration and proliferation, whilst treatment with a FURIN inhibitor reduces the effect of MeCP2 on monocyte migration and proliferation.

## Methods

### Generation of isogenic cell lines

THP1 monocytes were used as parental cells to generate isogenic cell lines with A/A or C/C genotypes at the rs17514846 position with CRISPR (clustered regularly interspaced short palindromic repeats)^5^. In brief, two gRNAs (guideRNA; 5′-GTCTGTGGGGGTCTCATTTTC-3′, and 5′-GCCCACATCCTCTGTTAAATG-3′, respectively) were separately cloned into the gRNA/Cas9 expression vector containing a green fluorescent protein reporter (p-UG-gRNA-EFS-Cas9-T2A-EGFP-WPRE). Additionally, a homology-directed repair donor vector was prepared by inserting 5’ and 3’ homology aims either side of the to-be-edited sites and subsequently subjected to site-directed mutagenesis to generate a vector containing the A allele at rs17514846. A PGK-Neo-PolyA cassette was also inserted in between the 5’ and 3’ arms in these vectors to provide a means for selection of edited THP-1 cells after transduction. THP1 cells, homozygous for the C allele of rs17514846, were transduced with the lentivirus particles containing the gRNAs and the A allele donor vector and incubated for 96 hours, followed by fluorescence-activated cell sorting. Sorted cells were cultured under G418 selection to select for successfully edited cells. The selected cells were screened via polymerase chain reaction (PCR) and Sanger sequencing to confirm successful editing of rs17514846 from the parental C/C genotype to A/A.

### Cell culture and 5-aza-2-deoxycytidine treatment

THP1 cells and the isogenic cell lines described above were cultured in RPMI 1640 medium supplemented with 10% fetal bovine serum and 1% penicillin-streptomycin at 37°C with 5% CO_2_. In some experiments, cells were first incubated with either 10µM 5-aza-2’-deoxycytidine (dissolved in dimethylsulfoxide) or the solvent only, for 72 hours.

### Quantitative reverse transcriptase polymerase chain reaction

Total RNA from isogenic THP1 cells was extracted with the use of Trizol reagent (Takara, 9109), reverse-transcribed into cDNA with a PrimeScript RT Reagent Kit (Takara, RR047A), and subjected to quantitative real-time polymerase chain reaction with a TB Green Premix Ex Taq II kit (Takara, RR820A). The PCR program consisted of 95°C for 1 minute, and then 40 cycles of 95°C for 15 seconds, 60°C for 30 seconds, 72°C for 30 seconds. The PCR primers were designed with the use of Primer-BLAST^11^ (https://www.ncbi.nlm.nih.gov/tools/primer-blast/index.cgi?ORGANISM=9606&INPUT_SEQUENCE=NM_001289823.2&LINK_LOC=nuccore). The sequences of the primers for amplifying *FURIN* transcript isoform 1&2 (NM_002569.4 and NM_001382622.1), *FURIN* transcript isoform 3 (NM_001289823.2), *FURIN* transcript isoform 4 (NM_001289824.2), an exon shared by the different *FURIN* isoforms, and the reference housekeeping gene *ACTB*, respectively, are shown in Supplementary Table I. *FURIN* expression levels were calculated with the use of the 2^-ΔΔCt^ method.

### Pyrosequencing

Genomic DNA was subjected to bisulfite modification and purification with the EpiJET bisulfite conversion kit (ThermoFisher, K1461), according to the manufacturer’s protocol. Briefly, 500ng DNA in 20µl was added into a 120µl modification reagent solution, mixed thoroughly and incubated at 98°C for 10 minutes and then 60°C for 4 hours. The modified DNA was purified, subjected to de-sulfonation at room temperature for 20 minutes, washed, and dissolved in 40µl elution buffer. The recovered DNA was used as a template in methylation-specific polymerase chain reaction (PCR) to amplify the target DNA segment, with biotin-labeled methylation-specific primers designed with Pyromark Assay Design Software 2.0 (Qiagen). The primer sequences are shown in Supplemental Table S1. The PCR products were mixed with magnetic beads coated with streptavidin and isolated with a Pyromark Q24 Vacuum Workstation (Qiagen), followed by pyrosequencing with a PyroMark Q24 instrument (Qiagen). The sequencing signals were analyzed with the use of Pyromark Q24 2.0.6 software (Qiagen).

### Bioinformatics analysis

PROMO^12,13^ (https://alggen.lsi.upc.es/cgi-bin/promo_v3/promo/promoinit.cgi?dirDB=TF_8.3) and TRANSFAC (http://gene-regulation.com/pub/databases.html) were used to search for transcription factor binding motifs in the DNA sequence at and surrounding the rs17514846 site. Subsequently, the implicated transcription factor was searched in the JASPAR^14^ (https://jaspar.genereg.net/) and Cistrom DB^15,16^ (http://cistrome.org/db/#/) databases to further check that its binding motif matched the sequence containing the rs17514846 site.

### Electrophoretic mobility super-shift assay

Nuclear proteins were extracted from THP1 cells with NE-PER Nuclear and Cytoplasmic Extraction Reagents (ThermoFisher, 78833). Biotin-labeled double-stranded oligonucleotide probes corresponding to the DNA sequences containing and surrounding the rs17514846 site (5’-AGTTGCGCCTGA[C/A]GCCTGCTTTCTT-3’), of either the C or A allele, were individually incubated with or without nuclear protein extracts, in the presence or absence of unlabeled C allele probe or unlabeled A allele probe or a unlabeled non-specific oligonucleotide, for 20 minutes at 4°C. For super-shift assay, the nuclear protein extracts were firstly incubated with a rabbit anti-human MeCP2 antibody or an equivalent amount of isotype rabbit IgG antibody for 30 minutes at 4°C, followed by an incubation with the biotin-labeled C allele probe for 20 minutes at 4°C. The mixes were electrophoresed on a 6.5% non-denaturing polyacrylamide gel and subsequently transferred onto a nylon membrane. The membrane was incubated with streptavidin-conjugated horseradish peroxidase and then with reagents of the Chemiluminescent Nucleic Acid Detection Module Kit (ThermoFisher, 89880). The membrane was then scanned with the use of a ChemiDoc XRS+ System (Bio-Rad).

### Chromatin immunoprecipitation and qPCR

Chromatin immunoprecipitation was performed with the use of a ChIP Kit (Abcam, ab270816). Briefly, THP1 cells were treated with 1% formaldehyde for 10 minutes to induce protein crosslinking. The crosslinking reaction was terminated by adding glycine. Thereafter, the cells were washed with ice-cold phosphate buffered saline and then lysed on ice with lysis buffers of the ChIP kit. The chromatin DNA was sheared into fragments of between 200 and 1,200 base pairs in length by sonication (Covaris). The chromatin DNA-protein complexes were incubated with an anti-MeCP2 antibody (Cell Signaling Technology, 3456) or an isotype control antibody (Abcam, ab37415) at 4°C overnight. The immunocomplexes were precipitated with protein A-coated magnetic beads. DNA in the immunocomplexes was then isolated and subjected to a quantitative PCR analysis of the *FURIN* gene, standardized against input DNA sample, using the 2^-ΔΔCt^ method. The sequences of the PCR primers are shown in Supplemental Table S1. The results are expressed as the fold difference by comparing the quantitative PCR values from the samples pulled down by the anti-MeCP2 antibody versus the quantitative PCR values from the samples pulled down by the isotype control antibody.

### Transfection of small interference RNA

Small interference RNA (siRNA) targeting *MeCP2* and a negative control siRNA were synthesized by GenePharma (Shanghai). The sequences of the *MeCP2* siRNA were 5′-GGAAAGGACUGACCUGUUUU-3′ and 5′-AACAGGUCUUCAGUCCUUUCCUU-3′. The sequences of the negative control siRNA were 5’-UUCUCCGAACGUGUCACGUTT-3’ and 5’-ACGUGACACGUUCGGAGAATT-3’. THP1 cells were transfected with either the *MeCP2* siRNA or control siRNA, with the use of Lipofectamine RNAiMAX Transfection Reagent (Invitrogen, 13778), and subjected to downstream experiments at 48 hours or 72 hours post transfection.

### Western blotting analysis

Protein extracts were prepared from THP1 cells and isogenic cells of the rs17514846 C/C genotype and A/A genotype, respectively. The protein extracts were subjected to denaturing polyacrylamide gel electrophoresis and then transferred onto a polyvinylidene difluoride membrane. The membrane was incubated with either a rabbit anti-FURIN antibody (Abcam, ab183495), a rabbit anti-MeCP2 antibody (Cell Signaling Technology, 3456), or a rabbit anti-β-actin antibody (Sangon Biotech, D110001) at 4°C overnight, washed and then incubated with a fluorescein-labeled goat anti-rabbit secondary antibody (LI-COR Biosciences, 926-32211). The protein bands were imaged with the use of a ChemiDoc XRS+ System (Bio-Rad) and analyzed with ImageJ software.

### Cell migration assay

THP1 cells were transfected with either the *MeCP2* siRNA or negative control siRNA for 48 hours and then incubated with 2.5 µM CellTracker Green CMFDA (5-chloromethylfluorescein diacetate) (Invitrogen, C7025) for 30 minutes. Thereafter, cells were washed, suspended in RPMI 1640 medium without serum, and seeded into the upper chamber of trans-wells on a 96-well plate with 7×10^4^ cells per well. RPMI 1640 medium complemented with 10% fetal calf serum was added into the lower chamber of each well. In some assays, the FURIN inhibitor decanoyl-RVKR-CMK (GLPBIO, GC15108) was additionally added to both the upper and lower chambers, at a final concentration of 10 µM. Cells were then incubated in a 5% CO_2_ atmosphere at 37 °C for 24 hours. Cells that had migrated onto the underside of the upper chamber were observed under a fluorescence microscope, and cells at 7 randomly selected areas were photographed and counted. The average number of cells in the 7 areas is presented.

### Cell proliferation assay

THP1 cells were transfected with either the *MeCP2* siRNA or negative control siRNA for 48 hours and then seeded on a 96-well plate with 1×10^4^ cells per well. In some assays, cells were incubated with the FURIN inhibitor decanoyl-RVKR-CMK (GLPBIO, GC15108) at a final concentration of 10 µM for 24 hours. Cells were then subjected to a proliferation assay with the use of Cell Counting Kit-8 (MedChemExpress, HY-K0301).

### Cell apoptosis assay

THP1 cells were transfected with either the *MeCP2* siRNA or negative control siRNA for 48 hours and then seeded on a 6-well plate with 3.5×10^5^ cells per well. In some assays, cells were incubated with the FURIN inhibitor decanoyl-RVKR-CMK (GLPBIO, GC15108) at a final concentration of 10 µM for 24 hours. Cells were then subjected to an apoptosis assay with the use of FITC Annexin V Apoptosis Detection Kit I (BD Pharmingen, 556547).

### Statistical analyses

Mann-Whitney test was used to ascertain differences between experimental groups in the *FURIN* expression level, fold enrichment of DNA in the chromatin immunoprecipitation-qPCR analysis, standardized Western blotting band intensity, and the rates of cell migration, proliferation, and apoptosis, respectively. A *P*<0.05 in a 2-tailed test was considered statistically significant.

## Results

### Allelic effect of rs17514846 on FURIN expression

To investigate the influence of rs17514846 on *FURIN* expression level, we performed a quantitative reverse transcriptase polymorphism chain reaction (qRT-PCR) analysis of the *FURIN* transcript isoforms^17^ in two isogenic cell lines (derived from THP1 monocytic cells) that were genetically identical, apart from the rs17514846 position where one of these isogenic lines was homozygous for the C allele whilst the other isogenic line was homozygous for the A allele. The analysis showed that isogenic cells of the rs17514846 A allele had higher expression of *FURIN*, particularly transcript isoforms 1&2 (NM_002569.4 and NM_001382622.1) and isoform 4 (NM_001289824.2), than isogenic cells of the C allele (Figure 1).

**Figure 1.**
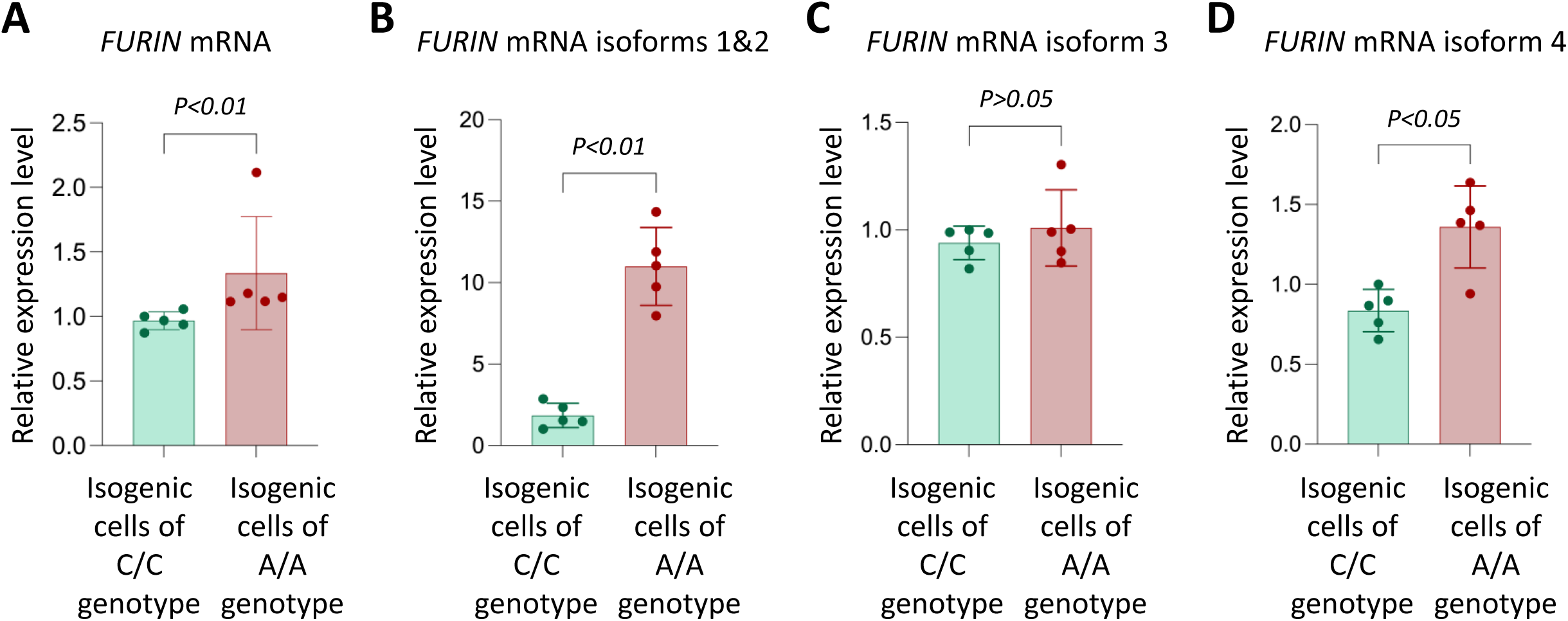
Monocytic cells of rs17514846 A/A genotype express higher levels of *FURIN* than monocytic cells of the C/C genotype. Results of qRT-PCR analysis of *FURIN* in isogenic monocytic cells of either the rs17514846 C/C or A/A genotype. Data shown are mean (± standard deviation) of *FURIN* relative expression levels standardized against the expression levels of the reference housekeeping gene *ACTB; P* values from Mann-Whitney test; n=5 in each group.

### Allelic effect of rs17514846 on DNA methylation

The rs17514846 SNP resides in a CpG site and the C to A change abolishes this CpG site, namely, the C allele (TTGCGCCTGA<u>C</u>GCCTGCTTTC), but not the A allele (TTGCGCCTGA<u>A</u>GCCTGCTTTC), comprises part of the CpG motif. Therefore, we wondered whether this CpG site was subjected to methylation. A pyrosequencing methylation analysis of THP1 monocytic cells (which were of the rs17514846 C/C genotype) showed a high level of 5-methylcytosine (5-mC) at this CpG site (Figure 2). Several other CpG motifs nearby were also found to be methylated (Figure 2).

**Figure 2.**
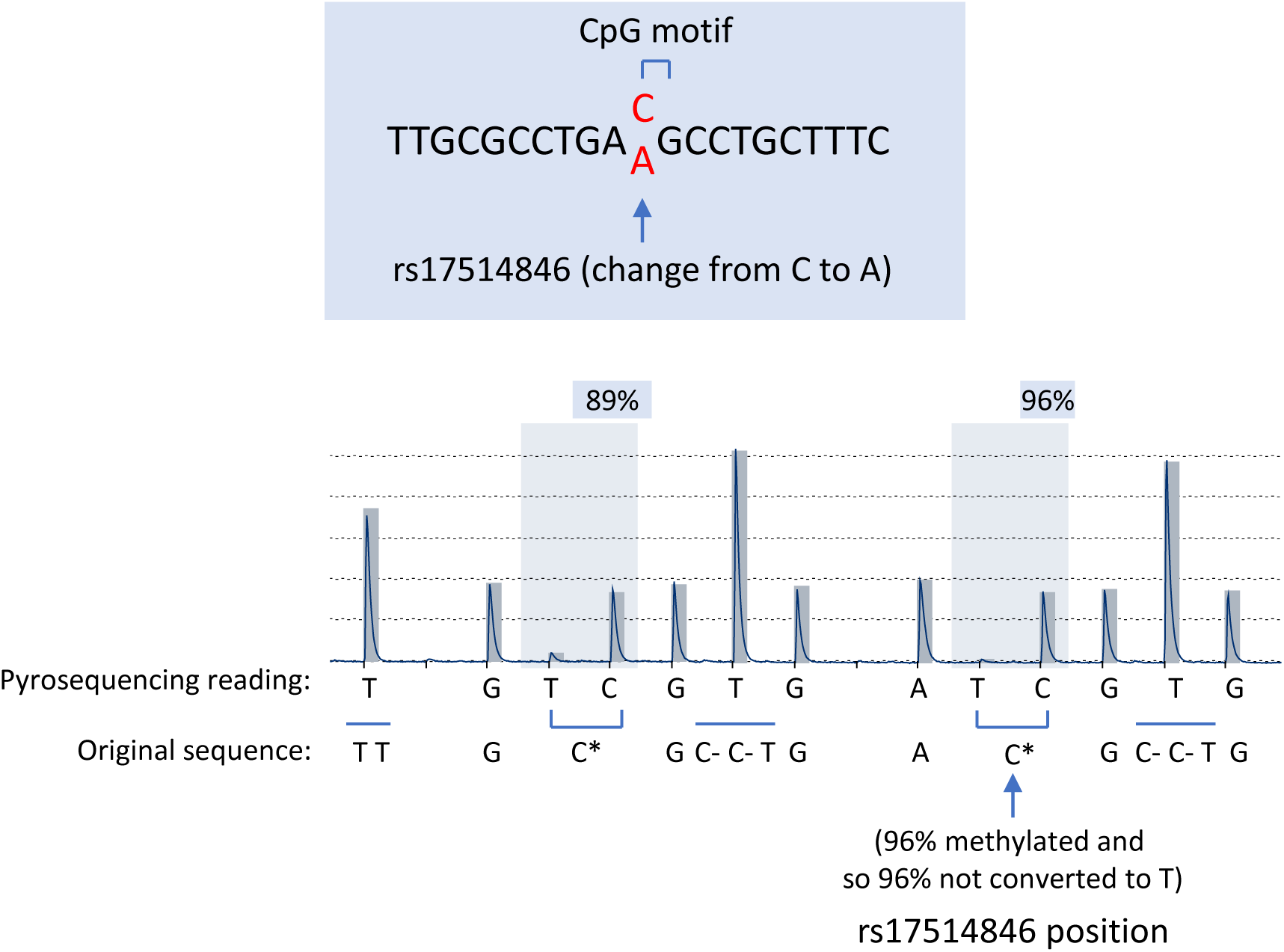
rs17514846 abolishes a methylated CpG motif. Upper panel: the rs17514846 resides in a CpG site and the C to A change abolishes this CpG site, namely, the C allele (TTGCGCCTGA<u>C</u>GCCTGCTTTC), but not the A allele (TTGCGCCTGA<u>A</u>GCCTGCTTTC), contains this CpG motif. Lower panel: a pyrosequencing methylation analysis of THP1 monocytic cells showed that in the rs17514846 C allele, the cytosine in this CpG motif was methylated. C* denotes methylated cytosine that was not converted to T in pyrosequencing; C-indicates unmethylated cytosine that was converted to T in pyrosequencing.

### DNA methylation inhibition increased FURIN expression

Since rs17514846 resides in a methylated CpG site, we sought to investigate if DNA methylation at this site influenced *FURIN* expression. We found that incubating THP1 monocytic cells with a DNA methylation inhibitor, 5-aza-2’-deoxycytidine, attenuated methylation of the cytosine at the rs17514846 site (Figure 3A). Furthermore, a qRT-PCR analysis showed that treatment with this DNA methylation inhibitor increased *FURIN* expression in isogenic cells of the rs17514846 C/C genotype, particularly, *FURIN* transcript isoforms 1&2 and isoform 4 (Figure 3B). In contrast, in isogenic cells of the A/A genotype, there were no significant changes in *FURIN* expression following treatment with the DNA methylation inhibitor (Figure 3B).

**Figure 3.**
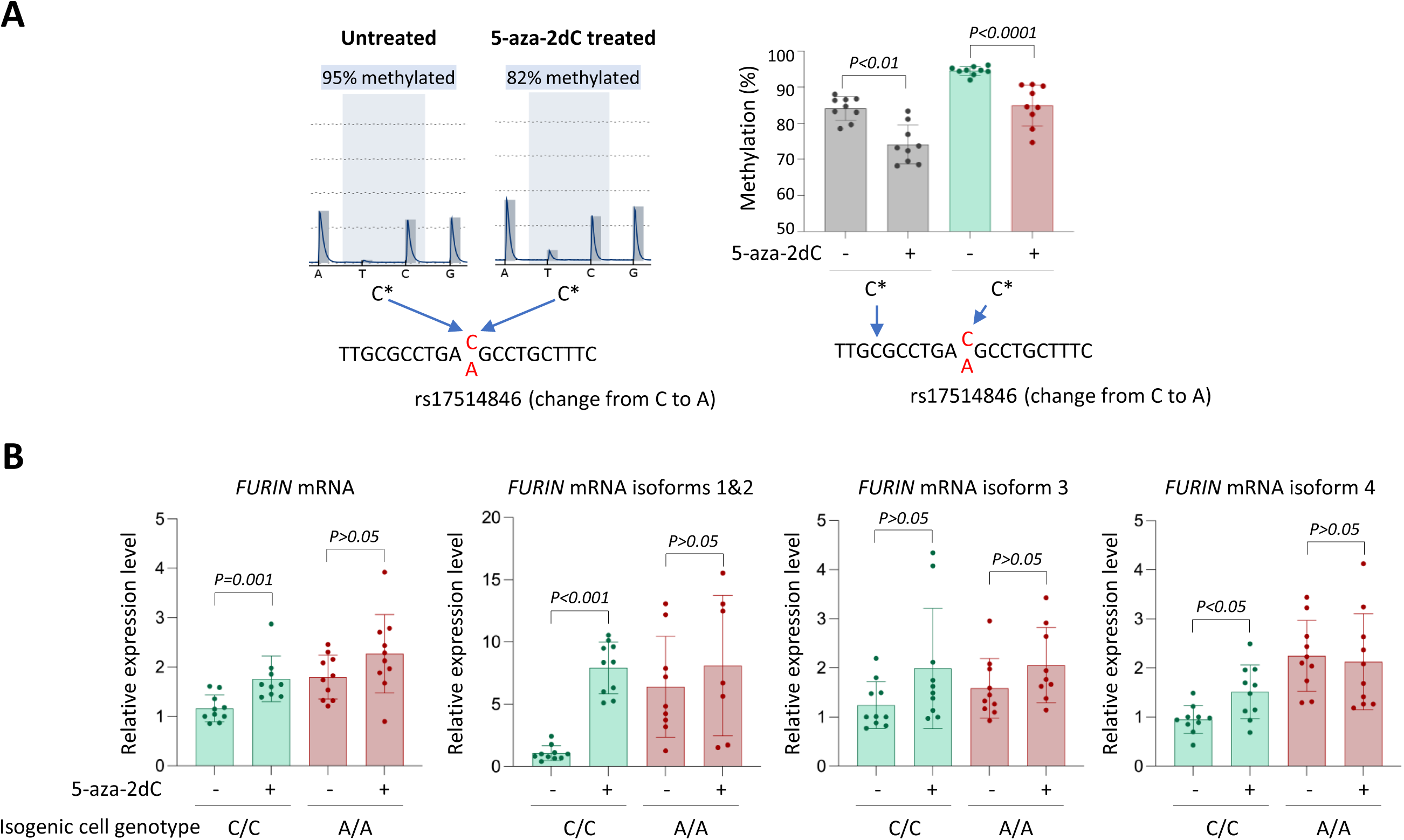
DNA methylation inhibition increases *FURIN* expression in isogenic monocytic cells of the rs17514846 C/C genotype. **A**. Pyrosequencing analysis of THP1 monocytic cells (of the rs17514846 C/C genotype) treated with the DNA methylation inhibitor 5-aza-2-deoxycytidine (5-aza-2dC, 10 µM) for 24 hours. Left: Average representative pyrograms at the rs17514846 position; Right: mean percentage (± standard deviation) of methylated cytosine at the respectively indicated position; *P* values from Mann Whitney test; n=9 in each group. **B**. *FURIN* RT-PCR analysis of isogenic monocytic cells of either the rs17514846 C/C or A/A genotype, treated with 5-aza-2dC (10 µM) for 72 hours. Data shown are mean (± standard deviation) of *FURIN* relative expression levels standardized against the expression levels of the reference housekeeping gene *ACTB*; *P* values from Mann-Whitney test; n=7-10 in each group.

### Allele-differential binding of the transcription factor MeCP2

Since the rs17514846 CpG site in the C allele is methylated and this CpG motif is abolished in the A allele, we wondered whether this would affect transcription factor binding to the DNA in this region. A bioinformatic analysis indicated a potential interaction of the rs17514846 site with the transcription factor MeCP2 that often functions as a gene repressor by binding to DNA with methylated CpG sites^9,10^.

In support of this hypothesis, an electrophoretic mobility super-shift assay using a probe corresponding to the DNA sequence at and surrounding rs17514846 with the C allele, together with THP1 monocytic cell nuclear protein extracts, detected DNA-protein complex bands (Figure 4A, lane 4) and showed that adding an anti-MeCP2 antibody in the assay resulted in an reduction in the intensity of one of these bands (Figure 4A, lanes 8 and 9, lower band) and a super-shift of the other band (Figure 4A, lanes 8 and 9, upper band), indicating binding of MeCP2 to the rs17514846 site of the C allele. In contrast, the DNA-protein complex bands were barely detectable using a probe corresponding to the A allele (Figure 4A, lane 2).

**Figure 4.**
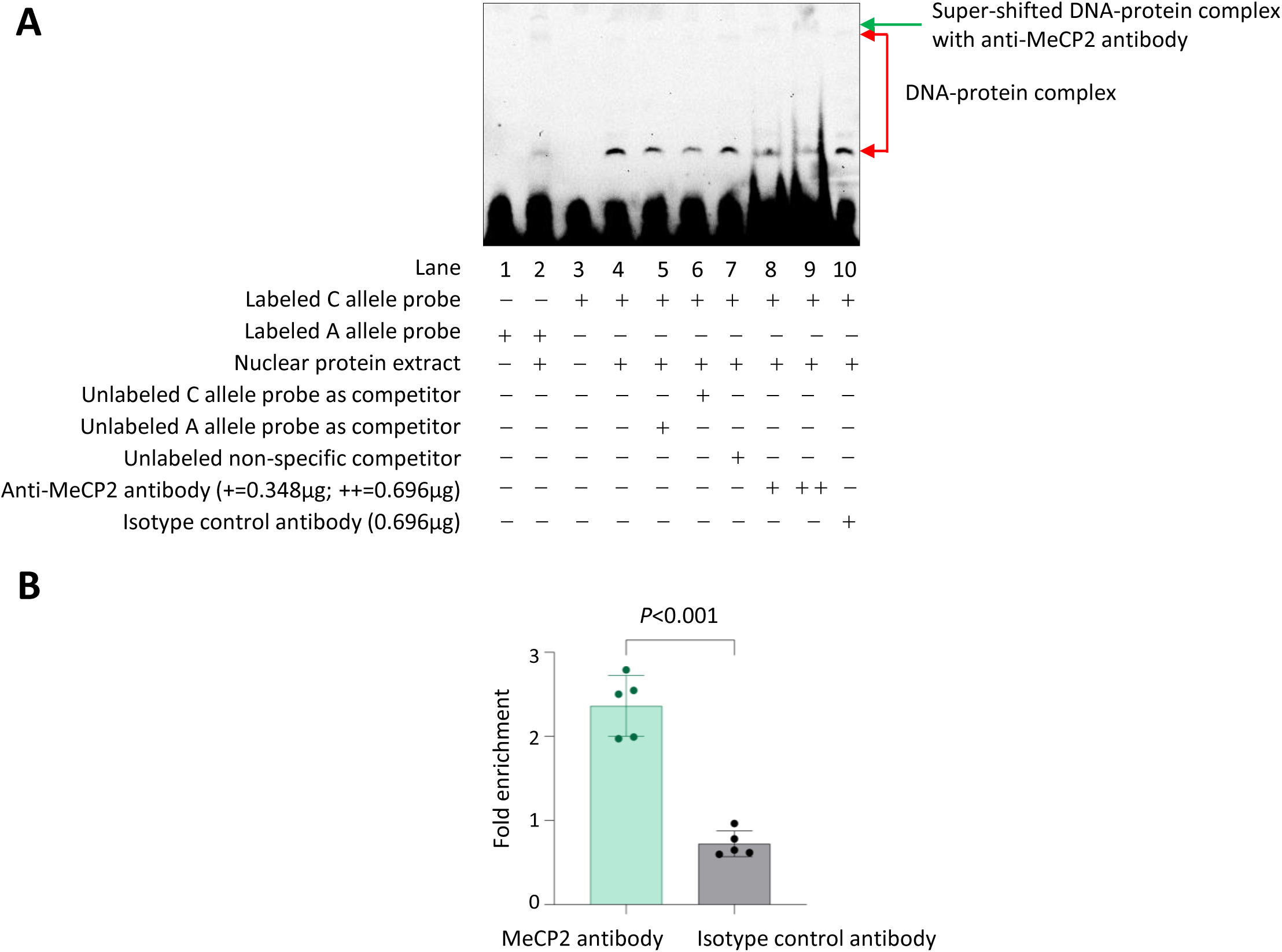
Preferential binding of MeCP2 to the rs17514846 C allele. **A**. An average representative image of electrophoretic mobility super-shift assay with probes corresponding to either the rs17514846 C or A allele, protein extracts from THP1 monocytic cells, an anti-MeCP2 antibody, and an isotype control antibody. **B**. Results of chromatin immunoprecipitation analysis of MeCP2 in THP1 cells. Data shown are mean (± standard deviation) of enrichment of the DNA sequence at and around the rs17514846 site in immunoprecipitated DNA sample as compared to input DNA sample, determined by quantitative PCR analysis. *P* values are from Mann-Whitney test; n=5 in each group.

In further support, a chromatin immunoprecipitation assay of THP1 monocytic cells (which were of the rs17514846 C/C genotype) detected an enrichment of the DNA sequence containing the rs17514846 site in chromatin precipitates pulled down by an anti-MeCP2 antibody, compared with chromatin precipitates with an isotype control antibody (Figure 4B).

### Knockdown of MeCP2 increased FURIN expression

Since MeCP2 often functions as a gene transcription repressor^9,10^, we investigated if knockdown of MeCP2 in cells would alter the FURIN expression level. The analysis showed that siRNA-mediated knockdown of MeCP2 increased FURIN expression in THP1 monocytic cells (which were of the rs17514846 C/C genotype) (Figure 5), suggesting that MeCP2 inhibits FURIN expression.

**Figure 5.**
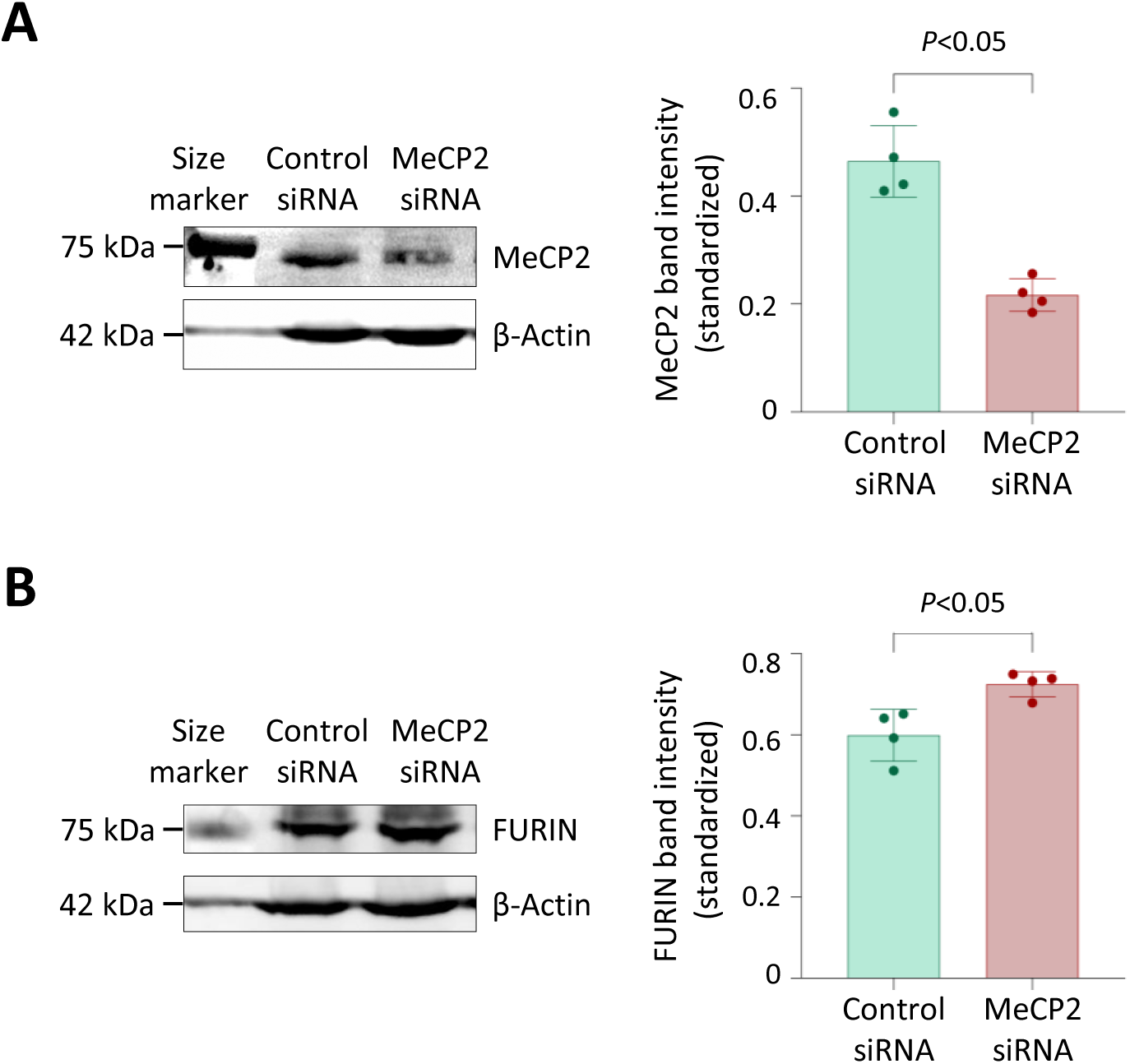
MeCP2 knockdown increases FURIN expression. Results of Western blotting analyses of MeCP2 (**A**) and FURIN (**B**) in THP1 monocytic cells transfected with either a *MeCP2* siRNA or negative control siRNA for 72 hours. Left: Average representative images of Western blotting images; Right: mean (± standard deviation) of MeCP2 band intensity (**A**) and FURIN band intensity (**B**), standardized against band intensity of the reference housekeeping protein β-actin. *P* values are from Mann-Whitney test; n=4 in each group.

### MeCP2 Knockdown increased monocyte migration and proliferation, and these effects were attenuated by a FURIN inhibitor

Previous studies have shown that FURIN promotes monocyte migration and proliferation^5,7^. Therefore, we investigated the possibility that the inhibitory effect of MeCP2 on FURIN expression has a bearing on monocyte migration and proliferation. We found that MeCP2 knockdown in THP1 monocytic cells increased their migration and proliferation, and this effect was diminished by a FURIN inhibitor (decanoyl-RVKR-CMK) (Figure 6A & 6B), suggesting that inhibition of FURIN expression by MeCP2 reduces monocyte migration and proliferation. MeCP2 knockdown did not affect the rate of monocyte apoptosis (Figure 6C).

**Figure 6.**
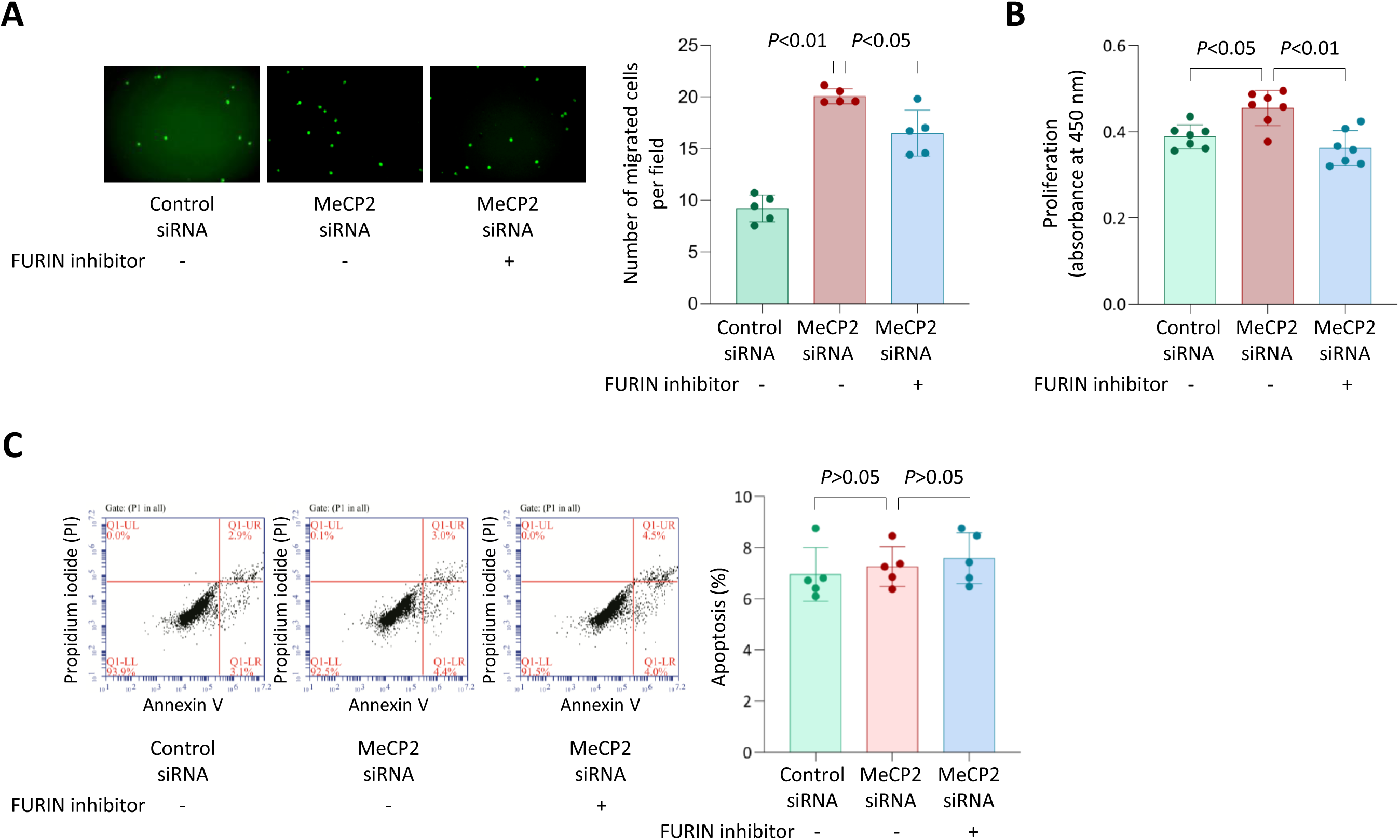
MeCP2 knockdown increases monocyte proliferation and migration, and these effects are attenuated by a FURIN inhibitor. Results of migration (**A**), proliferation (**B**) and apoptosis (**C**) assays of THP1 monocytic cells transfected with either a *MeCP2* siRNA or negative control siRNA for 48 hours, with or without treatment with the FURIN inhibitor decanoyl-RVKR-CMK for 24 hours. Data shown are mean (± standard deviation) values, *P* values from Mann-Whitney test; n=5 (**A** and **C**) or n=7 (**B**) in each group.

## Discussion

Genome-wide association studies have identified many genomic loci associated with CAD susceptibility^18,19^. However, the underlying biological mechanisms are still unclear for the majority of these loci, which hinders translation of the genetic discoveries into new treatments. Many of the identified CAD susceptibility loci are not associated with the classic risk factors such as hypercholesterolemia and hypertension,^1,20,21^ suggesting that they do not act through the traditional pathways and are thus not addressed by current treatments which primarily focus on lowering cholesterol levels and controlling blood pressure. Hence, there is a need to identify novel therapeutic targets for the development of new treatments that complement current strategies that target conventional risk factors.

rs17514846 has been reported as the lead CAD-associated SNP at the 15q26.1 locus in genome-wide association studies^1^ and this variant has been shown to influence FURIN expression in monocytes^5^. Our present study reveals a mechanism through which rs17514846 modulates FURIN expression. Our study shows that rs17514846 resides in a CpG motif in the presence of the C allele, which is methylated in monocytes. The C to A substitution producing the A allele of rs17514846 abolishes this methylated CpG motif. Our study reveals that the C allele interacts with MeCP2, a transcription factor known to be capable of binding to methylated DNA and act as a transcription repressor^9,10^. Furthermore, our study demonstrates that inhibition of DNA methylation with the use of 5-aza-2’-deoxycytidine or knockdown of MeCP2 can increase FURIN expression in monocytes. Taken together, these results suggest that the C to A substitution of rs17514846 abolishes DNA methylation at this site and thereby diminishes MeCP2 binding to this region, consequently leading to increased FURIN expression.

Previous studies have shown that *FURIN* expression is under the regulation of several transcription factors interacting with their respective binding sites in the *FURIN* regulatory regions, including C/EBP-β (CCAAT enhancer binding protein β)^17^, HIF1 (hypoxia-inducible actor-1)^22^, GATA1 (GATA binding protein 1)^23^, SMAD2/4 (SMAD family member 2 and SMAD family member 4)^24^, and STAT4 (signal transducer and activator of transcription 4)^25^. Our present study uncovers that *FURIN* expression is regulated also by the transcription factor MeCP2 that binds to the rs17514846 site in intron 1 of the *FURIN* gene. Whereas C/EBP-β, HIF1, GATA1 SMAD2/4 and STAT4 enhance *FURIN* expression as shown in previous studies^17,22–25^, MeCP2 displayed an inhibitory effect on *FURIN* expression in our study. Taken together, the findings from the previously reported studies^17,22–25^ and our present work indicate that *FURIN* expression is subjected to both positive and negative regulation.

FURIN is a potential therapeutic target associated with many diseases including atherosclerosis^7^ and cancers^26^, and therefore there has been considerable research into the development of FURIN inhibitors^27^. The findings of our present study indicate a possible new approach for targeting *FURIN*, namely, inhibiting *FURIN* expression by inducing/increasing DNA methylation in the *FURIN* gene. There are emerging methodologies that can potentially be used to achieve this. For example, it has been demonstrated that an inactive Cas9 fused with an engineered DNA methyltransferase can induce locus-specific DNA methylation without affecting global methylation^28^. Thus, controlling *FURIN* expression by inducing targeted DNA methylation offers another strategy for FURIN targeting, which warrants further investigation.

Our study demonstrates that a genetic variant can modulate gene expression through an epigenetic mechanism. As such a mechanism might operate at other loci as well, a possible genetics-epigenetics interaction would be worth considering in mechanistic studies of other disease-associated loci.

In summary, the results of our study provide a novel mechanistic understanding of the effect of the CAD-associated variant rs17514846 on *FURIN* expression and monocyte migration/proliferation. The finding that DNA methylation suppresses *FURIN* expression suggests that inducing targeted DNA methylation at the *FURIN* gene could be considered as a possible approach for FURIN inhibition in drug development.

## Highlights

- The coronary artery disease predisposing genetic variant rs17514846 abolishes a methylated CpG motif in the *FURIN* gene.
- The rs17514846 C allele interacts with the transcription factor MeCP2 that is known to often function as a gene repressor by binding to methylated CpG sites.
- Treatment with a DNA methylation inhibitor increases FURIN expression.
- MeCP2 attenuation increases FURIN expression.
- MeCP2 attenuation increases monocyte migration and proliferation, whereas a FURIN inhibitor diminishes this effect.

